# Automated localization of calling birds with small passive acoustic arrays in complex soundscapes

**DOI:** 10.64898/2026.02.23.707497

**Authors:** Michael B. Eisen, Patrick O. Brown, Ana Sanz Matias

## Abstract

Accurately localizing vocalizing animals from passive acoustic recordings remains challenging in complex outdoor soundscapes. Although automated detection and classification of bird calls have advanced rapidly, reliable spatial localization has lagged, particularly for small arrays of autonomous recorders operating without manual intervention. Here we describe a fully automated pipeline for three-dimensional localization of bird vocalizations using distributed networks of 4 to 8 GPS-synchronized recorders deployed in heterogeneous forest-adjacent environments. Our framework integrates automated multi-recorder event matching, frequency-selective FFT-based cross-correlation for time-difference-of-arrival (TDOA) estimation, geometric cycle-consistency filtering to resolve ambiguous cross-correlation peaks, and nonlinear optimization of source location and effective sound speed. We applied this pipeline to recordings collected from October 2023 through May 2026 at three sites in southwest Arkansas. Of 6,037,004 non-redundant high-confidence bird detections across 258 species, 1,381,317 passed our criteria for potential localization, of which 1,055,021 (76.4%) spanning 174 species yielded positional estimates. The localizations exhibit strong concordance with independently known landscape features and species-specific habitat associations. These analyses indicate that small, practical arrays can recover ecologically meaningful spatial structure in complex soundscapes without manual curation. This preprint documents the current state of the system and its performance under realistic field conditions.

## Introduction

Passive acoustic monitoring (PAM) has transformed the study of birds and other vocal wildlife by enabling continuous, non-invasive recording across broad spatial and temporal scales (Aide et al., 2013; Gibb et al., 2019; Shonfield and Bayne, 2017; Sugai et al., 2019). Autonomous recording units (ARUs) can now operate for weeks or months at a time, generating detailed records of species presence, seasonal activity, and behavioral timing. The rapid development of machine learning approaches for automated detection and classification - exemplified by systems such as BirdNET (Kahl et al., 2021) - has further accelerated this transformation, allowing researchers to extract species-level information from vast acoustic datasets. As a result, PAM has become a central tool in biodiversity assessment and ecological monitoring.

Despite these advances, most acoustic monitoring remains spatially implicit. A recorder can indicate which species are vocalizing nearby and how frequently they call, but it does not directly reveal where individuals are located in space. For many ecological questions - including territory structure, habitat use, vertical stratification of activity, social interactions, movement behavior, and density estimation - spatial information is essential. Without localization, acoustic data provide presence but not position.

The principles of acoustic localization are well established. When a sound reaches multiple microphones at slightly different times, differences in arrival time can be used to infer the source location. Time-difference-of-arrival (TDOA) localization has a long history in acoustics and radar, where position is estimated from hyperbolic constraints derived from pairwise arrival-time differences (Smith and Abel, 1987). In ecological contexts, tools such as Sound Finder made these methods accessible and demonstrated that distributed microphone arrays can recover positions of vocalizing animals under favorable conditions (Wilson et al., 2014). Subsequent reviews have synthesized advances in acoustic localization and highlighted both its promise and its practical constraints, particularly with respect to synchronization, environmental variability, and automation (Rhinehart et al., 2020).

Three-dimensional localization of birds has also been demonstrated, including recovery of flight altitude using arrays designed for that purpose (Dutilleux et al., 2023; Stepanian et al., 2016). These studies showed that acoustic arrays can move beyond presence detection to reconstruct animal positions in space. However, accurate localization typically requires careful array geometry, precise time synchronization, and relatively clean acoustic conditions. Many prior systems relied on manual curation, specialized hardware, or larger engineered arrays.

Open-source toolkits such as OpenSoundscape (Lapp et al., 2023) now provide accessible implementations of TDOA-based localization built around classical hyperbolic solvers. These tools illustrate the growing availability of reproducible acoustic localization workflows. However, there remains a gap between technical feasibility and routine, fully automated field deployment using small, logistically practical arrays of autonomous recorders.

In real forest soundscapes, multiple birds often vocalize simultaneously. Wind, insects, and anthropogenic noise introduce additional complexity. Autonomous recorders can drift in timing over days or weeks, introducing small synchronization errors that become consequential when millisecond precision is required. When small numbers of recorders are available, redundancy is limited. Misalignments in time or incorrect association of overlapping calls can produce localization estimates that appear geometrically plausible but are in fact incorrect. Altitude estimation is especially sensitive to such errors in structurally complex habitats where vegetation alters sound propagation.

Here, we present a fully automated framework for three-dimensional localization of bird vocalizations using small distributed arrays deployed in heterogeneous forest environments. Our approach integrates automated multi-recorder event matching, frequency-selective FFT-based cross-correlation for TDOA estimation, explicit modeling of recorder clock drift, and nonlinear optimization of source location and effective sound speed (Cramer, 1993). Critically, we introduce a geometric cycle-consistency filtering step prior to localization. Rather than accepting a single peak from each cross-correlation function, we evaluate combinations of candidate peaks under additive triangle constraints across recorder triplets and select those that minimize cycle residuals. This filtering enforces TDOA consistency around cycles in the recorder graph, echoing related to cycle-consistency constraints in synchronization problems (Singer, 2011), and reduces sensitivity to spurious or ambiguous cross-correlation peaks.

Our goal is not merely to demonstrate that localization is possible, but to show that it can be achieved reliably and without manual intervention under realistic field conditions using modest hardware. Applied to multi-year datasets from three arrays of autonomous recorders at different locations in a field site in southwest Arkansas, the system processes over a million candidate events and produces spatially coherent localizations consistent with landscape structure and species-specific habitat associations.

This work makes three contributions. First, we demonstrate that small, logistically practical arrays (4 to 8 recorders) can achieve robust three-dimensional localization in complex soundscapes without manual curation. Second, we introduce a geometric cycle-consistency filtering strategy that resolves ambiguous cross-correlation peak combinations under combinatorial uncertainty. Third, we present a fully automated, high-throughput pipeline that integrates machine-learning-based detection, TDOA estimation, sound-speed calibration, and nonlinear optimization into a system capable of operating at ecological timescales.

By enabling spatially explicit inference from small autonomous arrays, this framework expands the ecological value of passive acoustic monitoring from presence detection toward quantitative mapping of animal position, movement, and vertical habitat use in complex forest environments.

## Localization Strategy

### Array design and synchronization

We deployed four distributed arrays of autonomous recorders (4 to 8 recorders per site over the course of the study period, with efforts made to maintain at least 4 active recorders at any given time) across a 1,000-acre field site in southwestern Arkansas. Solar BAR (Frontier Labs) recorders were spaced approximately 35 m apart. This spacing ensures that inter-recorder delays can be well-resolved at standard audio sampling rates. We also added one Song Meter SM4-TS Acoustic Recorder (Wildlife Acoustics) elevated approximately 7 m at each site for some of the time periods analyzed. Recorder positions were measured using an Emlid Reach RS2 GNSS receiver with differential corrections, yielding sub-meter positional accuracy.

All recorders were equipped with GPS receivers (pre-installed for Solar BAR and added for SM4) used to provide the timing precision to recordings needed to recover positions. For Solar BARs, each file is named with GPS-derived start and stop timestamps to microsecond precision, from which the recorder’s true sample rate over that file is recovered as the frame count divided by the elapsed GPS time. This calibration is essential rather than cosmetic: the recorders’ oscillators run approximately 24 ppm slow and differ from one another by 23 ppm (44098.53 to 44099.54 Hz against a nominal 44100 Hz), so two uncalibrated recorders accumulate one millisecond of relative timing error (about 0.35 m of path length) every 44 seconds. Applying the per-file rate removes this drift, and the residual file-to-file variation within a recorder is 3 ppm.

Elevated SM4-TS recorders are GPS-disciplined: recording begins within a fraction of a millisecond of GPS time, and the recorder continuously adjusts its sample rate during recording to maintain synchronization to within 1 ms. Files for which synchronization was acquired before recording and held throughout are flagged by the manufacturer in the filename, and only those files are used.

Each array operated autonomously for extended periods. Due to environmental and logistical factors (e.g., flooding, ant invasions, power interruptions, acts of hog), some recorders were intermittently offline; localization analyses were restricted to periods when at least four recorders were active at the relevant site.

### Detection and event selection

BirdNET detections were performed on synchronized, non-overlapping 3-second windows. For TDOA estimation, we analyzed a 5-second segment centered on the detected window (adding one second of padding on either side) to capture call energy overlapping window boundaries. Window start times were aligned across recorders to facilitate inter-recorder comparison.

For the analyses presented here, we considered only species known to occur at the site and retained high-confidence detections (BirdNET confidence ≥ 0.9). We required that a candidate event be detected on at least three recorders within the same 3-second window and that at least four recorders were active at that time.

We did not attempt to temporally isolate individual calls within the 3-second window. Preliminary attempts to refine call onset times using envelope-based, spectrogram-based, and template-based approaches did not consistently identify calls, nor did they reliably improve localization performance in complex soundscapes with overlapping signals. Instead, we estimate TDOAs using a 5-second segment centered on the 3-second detection window, providing padding to capture call energy that overlaps window boundaries, and rely on downstream geometric consistency filtering to identify the correct delay combination. While sub-window localization may further improve performance, we consider it a future direction.

### TDOA estimation

Time-difference-of-arrival (TDOA) localization has a long history in acoustics and radar, where position is estimated from pairwise arrival-time differences under geometric constraints (Smith and Abel, 1987). We estimated pairwise TDOAs using FFT-based generalized cross-correlation (Knapp and Carter, 1976). Rather than applying broadband GCC-PHAT weighting, which equalizes spectral power across frequencies, we implemented frequency-selective weighting derived from spectral profiles of focal species vocalizations, built from reference recordings in The Cornell Guide to Bird Sounds: United States and Canada (Cornell Lab of Ornithology, Macaulay Library). In preliminary analyses, broadband PHAT weighting amplified insect and anthropogenic noise, degrading peak definition in forest soundscapes. Frequency-selective weighting improved peak discrimination while preserving temporal resolution. Recorder pairs lacking well-defined peaks within the physically feasible delay range were excluded from further analysis.

### Geometric consistency filtering

Cross-correlation functions frequently contained multiple plausible peaks due to overlapping calls and internal call structure. Rather than selecting the largest peak per pair, we retained all candidate peaks above a signal-to-noise threshold within the physically feasible delay range, and exploited a geometric constraint to select from all possible combinations consisting of one candidate peak per pair, only those with consistent geometry.

Under an internally consistent sign convention and in the absence of measurement error, true TDOAs satisfy additive closure:

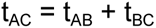

for all triplets of recorders A, B, C. In practice, measurement noise and reverberation introduce deviations from exact closure; our objective function penalizes these deviations rather than requiring exact equality.

We evaluated each combination of plausible peaks according to its total deviation from triangle closure across all recorder triplets. The optimal peak set minimized the summed squared closure residual across all recorder triplets.

The triangle consistency constraints used here are closely related to cycle-consistency conditions in graph-based synchronization problems, where pairwise measurements must agree around cycles in a measurement graph. Similar ideas appear in angular synchronization and related estimation problems (Singer, 2011). In our case, the cycles are length-3 recorder triplets and the constraints enforce additive consistency of TDOAs.

This cycle-consistency approach allows identification of mutually compatible TDOA sets even when the correct peak is not the largest individual cross-correlation maximum. This can be done efficiently. In practice, the number of candidate peaks per recorder pair was typically 2 to 5 after signal-to-noise filtering, resulting in manageable combinatorial search spaces. When peak counts were unusually large, we constrained candidate sets based on peak prominence and delay plausibility to prevent combinatorial explosion.

In many events, the optimal peak set corresponded to the largest cross-correlation peak for each recorder pair. However, in cases with overlapping calls or strong internal structure, smaller peaks were occasionally selected because they formed geometrically consistent combinations.

### Speed of sound estimation

Accurate localization requires knowledge of sound speed, which varies with temperature and humidity (Cramer, 1993). At our site, air temperature at the time of analyzed detections ranged from −13 °C to 42 °C, across which the speed of sound varies by 34 m·s⁻¹; 90% of analyzed calls fall within a narrower band of 17 m·s⁻¹ (336.0 to 352.7 m·s⁻¹).

We estimated the speed of sound for each event from the hourly 2 m air temperature and relative humidity at the site and call time, taken from the HRRR 3 km analysis via the

Open-Meteo historical-forecast archive, applying the temperature-dependent speed-of-sound relation of Cramer (1993) together with the humidity correction c = c_dry (1 + 0.165 x_w), where x_w is the water-vapor mole fraction.

This temperature- and humidity-derived value was used directly as the effective sound speed and was not refined during optimization. Allowing sound speed to vary per event renders the four-recorder geometry poorly conditioned, since a change in sound speed trades off against a radial shift in source position.

### Localization optimization

Source location was estimated via nonlinear least-squares minimization of the residual between observed and predicted TDOAs given recorder positions. We used nonlinear least-squares optimization (SciPy least_squares, trust-region reflective algorithm (Virtanen et al., 2020)). We required at least four recorders with valid TDOAs to solve the three-dimensional localization problem.

Pairs with inconsistent or noisy TDOAs were excluded prior to optimization. Solutions were accepted only if they satisfied thresholds for geometric closure deviation and overall residual error.

The TDOA cost surface is multimodal. Re-solving the same events from a naive initialization at the array centroid recovers the reported minimum for 56.0% of calls; the remainder converge to a solution a median of 61 m away whose residual is substantially larger and which would fail our 1 ms quality threshold in 99.7% of cases, against 33.5% for the reported solution. Peak selection and initialization therefore do substantive work, and the residual threshold reliably distinguishes the two outcomes. In a small minority of cases (3.2% of all calls) the alternative minimum has the lower residual, but these are events whose residual is of order 46 ms under either solution (far above our acceptance threshold) so they are excluded from the reported corpus regardless of which minimum is found.

## Results

We analyzed synchronized recordings collected from October 2023 through May 2026 across three field sites (Figure 1). Automated BirdNET-Analyzer processing identified 6,037,004 non-redundant high-confidence bird detections (confidence ≥ 0.9) across 258 species, drawn from a site species list of 300 expected bird species; 161 of these species were detected more than 100 times and account for 99.96% of all detections. Restricting to events detected on at least three recorders within the same 3-second window, and limiting to periods when at least four recorders were active, yielding 1,381,317 candidates for localization. Applying our localization pipeline produced position estimates for 1,055,021 of these events (76.4%), spanning 174 species; the remainder failed a required geometric constraint (an insufficient number of recorder triplets satisfying the triangle-closure criterion described below).

**Figure 1.**
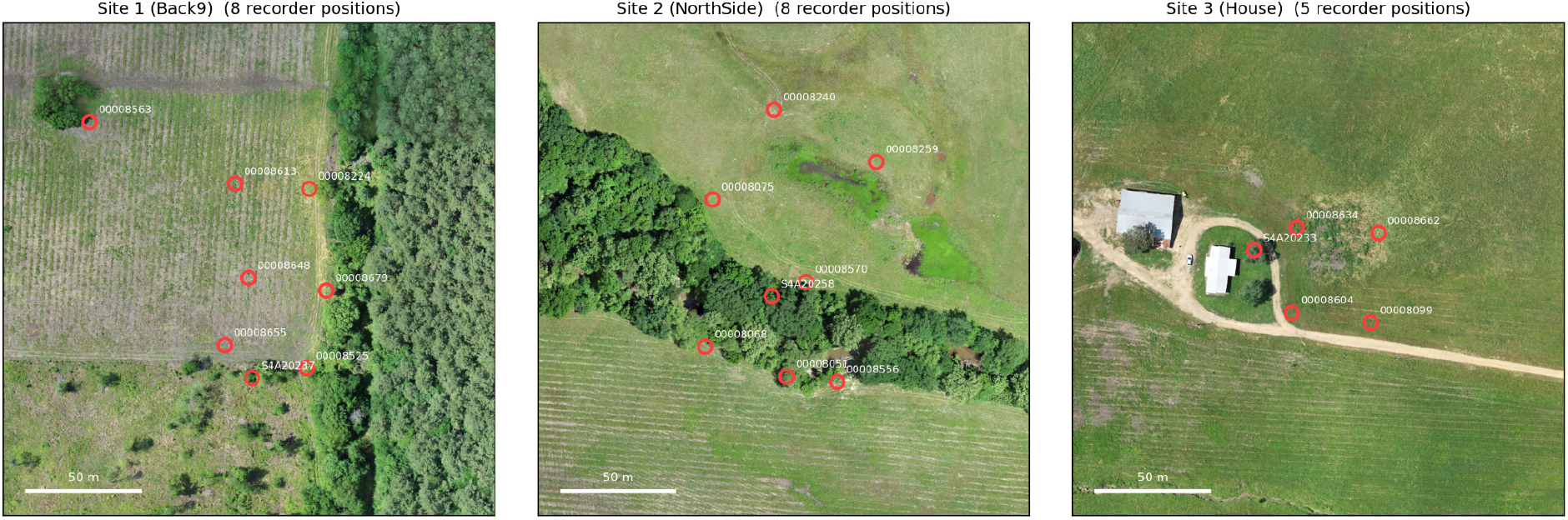
Spatial configuration of passive acoustic monitoring arrays at three field sites. Locations of GPS-synchronized autonomous recorders deployed at each site, shown on drone orthophotography acquired in May 2025. Red circles mark surveyed recorder positions, measured with an Emlid Reach RS2 GNSS receiver with differential corrections (sub-meter accuracy). Each panel spans the same ground area. Maximum within-array separations are 90 m (Site 1), 113 m (Site 2) and 54 m (Site 3). Counts give the number of distinct recorder positions occupied over the study period rather than the number operating simultaneously, which ranged from four to six per site: recorders were occasionally relocated or replaced, and each deployment is plotted separately. Song Meter SM4-TS recorders (prefix S4A) positioned at a relative elevation of 7m are included, although they were not continuously operating.

We assessed the overall performance of the algorithm in three ways: 1) By examining the distribution of mean residual TDOA errors after optimization (the difference between the observed TDOAs and those inferred from the localized position) and how different aspects of our methodology affects it, 2) By manual inspection of input spectrograms aligned based on predicted relative time of arrivals (TOA) of a call emitted from the predicted location, and 3) By examining the relationship between call localizations and spatially localized features surrounding each recorder array, especially trees and other perching sites. The first two metrics reflect model fit and convergence quality; the third provides the most direct, if indirect, evidence of spatial accuracy.

The median residual TDOA error across localizations is 0.89 ms (Figures 2A & B), corresponding to a mean path-length discrepancy of about 0.31 m. … Introducing random positional perturbations of 5 m and 10 m to recorder coordinates — each recorder displaced by exactly that distance in a uniformly random three-dimensional direction — substantially increased residual TDOA error, from 0.47 ms unperturbed to medians of 3.72 ms and 8.20 ms respectively, and displaced the resulting positions by medians of 12 m and 20 m horizontally.

**Figure 2.**
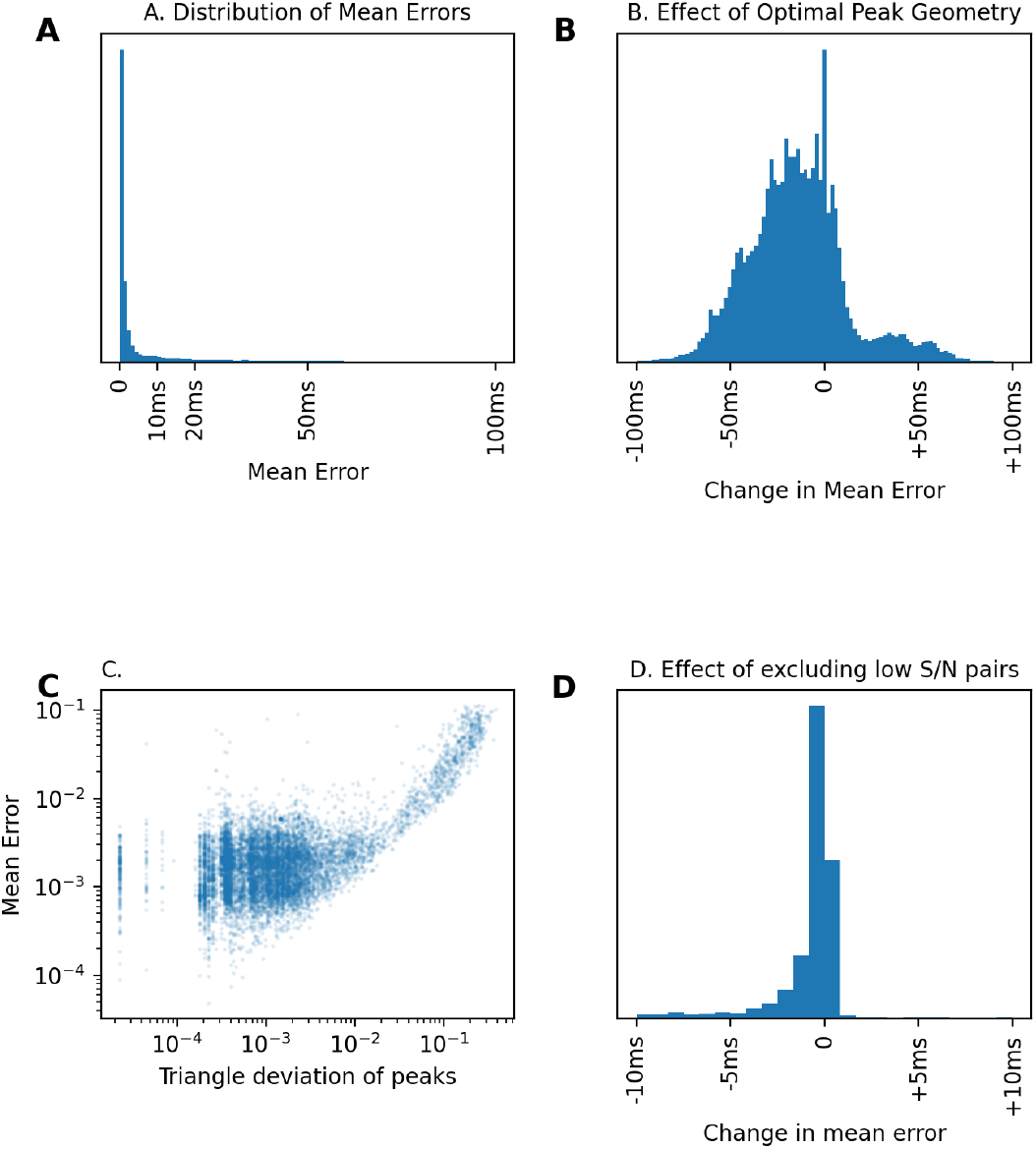
Internal geometric consistency and sensitivity to recorder position perturbation. (A) Distribution of mean residual TDOA error across all localized events (n = 1,055,021), defined as the difference between observed TDOAs and those predicted from the optimized source position at the temperature- and humidity-derived sound speed. The median error is 0.89 ms, equivalent to a path-length discrepancy of about 0.31 m, and 52.5% of events fall below 1 ms. (B) Change in mean residual TDOA error between the peak set selected by geometric consistency and the peak set formed by taking the largest cross-correlation peak for each pair, restricted to the events where the two differ. Negative values indicate lower error under geometric selection. (C) Relationship between geometry score and mean residual error, restricted to six-recorder events in which all 15 recorder pairs passed quality filtering and none were dropped (n = 10,153), so that differences reflect peak selection rather than differing numbers of constraints. (D) Change in mean residual TDOA error when recorder pairs with low signal-to-noise are excluded by a stricter cross-correlation quality cutoff (0.05 rather than the default 0.2). Negative values indicate lower error under the stricter cut. Only 7.7% of candidates (1,534 of 20,000 sampled) retained enough usable pairs to be re-localized at the stricter cutoff; the panel therefore shows how much the surviving events improved, not what the stricter cutoff would cost, which is the loss of roughly 92% of the data.

Inferred TOAs from the localization solution should temporally align the focal call across recorders. While the existence of a good alignment does not imply accurate localization (there are some geometric degeneracies in our setup, and there is some susceptibility to over-fitting), the failure to align calls across recorders represents localization failure. Although we do not yet quantify alignment quality algorithmically, visual inspection (Figure 3) indicates that misaligned events correspond to large residual errors (>10 ms). We also implemented a measure of temporal support for each recorder pair (Figure 3, top panels) to highlight when the triangle closure criteria selected peaks emerging from overlapping calls in different parts of the temporal window, another indicator of likely localization failure.

**Figure 3.**
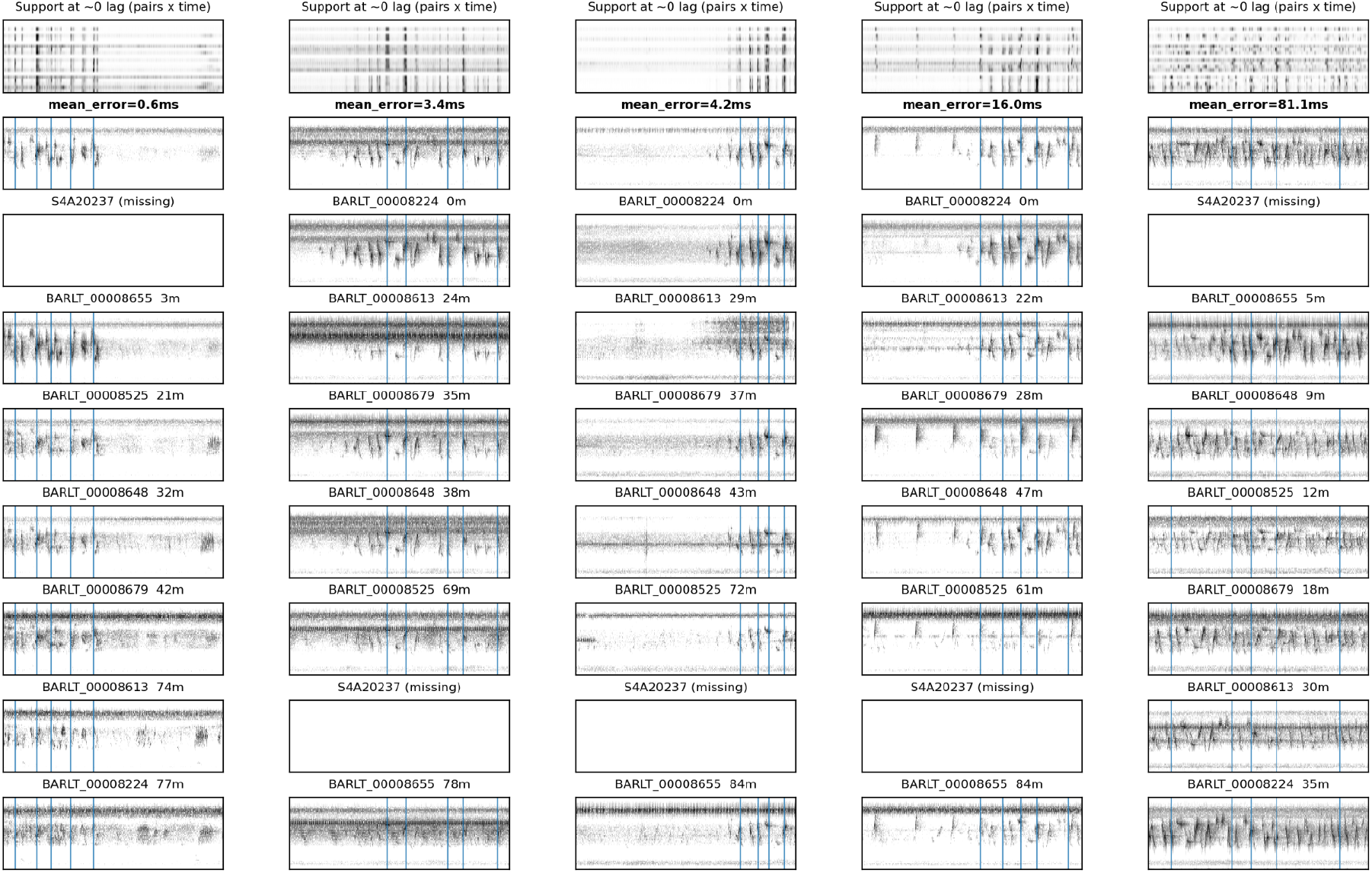
TOA-aligned spectrograms across a range of localization quality. Representative multi-recorder spectrograms aligned according to times of arrival inferred from the localization solution: each recorder’s audio is shifted by the propagation delay predicted from the estimated source position and sound speed, so that a correctly localized call becomes vertically aligned across the stack. Each column shows one event. Within a column, rows give the per-recorder spectrograms beneath a support map and the mean aligned spectrogram across recorders. Audio is band-weighted using the same species-specific frequency profile used for TDOA estimation, so the panels show what the cross-correlation operates on rather than the raw recording. Events are ordered by mean residual TDOA error. For well-localized events the focal call aligns across recorders. Misaligned examples correspond to events with elevated residual error, typically where overlapping vocalizations or low signal-to-noise left the correct cross-correlation peak ambiguous.

Although we have not yet systematically surveyed call locations independent of audio recordings to serve as ground truth, a wealth of observational data on the relevant species, and our experience from many months working at each site, suggests that the majority of bird calls are made from trees and other elevated perching locations. We therefore analyzed high quality calls (mean error less than 1 ms) against the physical features at each site (Figure 4), and the concordance with tree and other perching locations is qualitatively strong.

**Figure 4.**
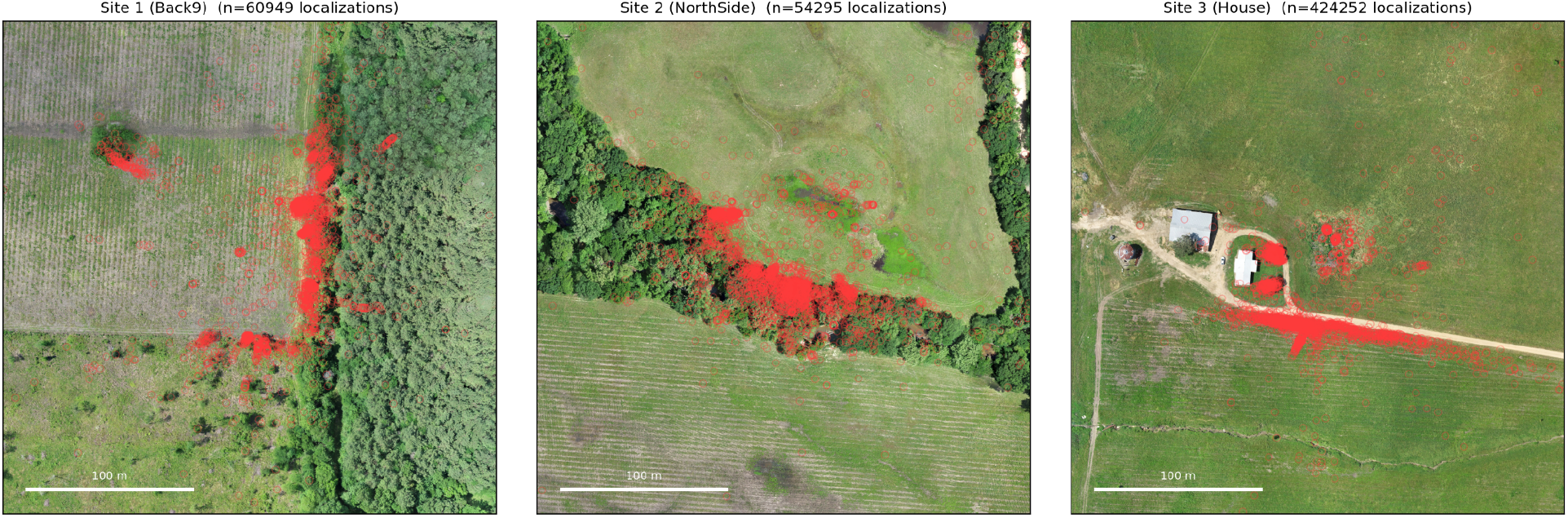
Spatial distribution of high-quality localizations relative to landscape structure. Localizations with mean residual TDOA error below 1 ms (n = 553,615 of 1,055,021) overlaid on drone orthophotography of each site. Each panel spans 300 m centered on the array centroid. Clusters of inferred call locations correspond to treelines, isolated trees, forest edges, powerlines and other elevated perching sites, and are largely absent from open ground. These spatial patterns are consistent with independent field observations and with expected species behavior, and constitute the principal external evidence that the inferred positions are accurate: the features they recover are meters wide and were not supplied to the algorithm in any form. Note that panels are plotted from a random subsample of at most 3,000 points each to avoid saturation, so plotted point density is not proportional to the per-panel counts given in the titles.

At site 1, where we have six recorders localized in a grid in a field adjacent to a pine forest and a perpendicular treeline, we see most of the localizations along the treeline to the east of the recorder array, in specific trees along the southern field border, and in a small isolated grove of trees to the northwest of the recorders. At site 2, which is situated on either side of a forested creek, with areas of flooding and vernal pools to the north, the localizations follow the treeline closely, with additional clusters at the seasonal pools. At site 3, which is adjacent to a house and barn, the localizations cluster in the two trees to the front (east) of the house, in an uncleared and slightly depressed thicket further to the east, and along the powerline that runs parallel to and on the south side of the road. Site 4 is a small copse of trees along a seasonal creek; we have collected less data here than at the other three sites, and defer a detailed spatial analysis to a later version of this manuscript alongside updated figures. The patterns at sites 1-3 correspond closely to our subjective impression of where most calls at each site originate.

Finally, we looked at the spatial localization of individual species (Figure 5), which again match their ecological expectations and our subjective observations. Indigo buntings (*Passerina cyanea*) are abundant at sites 1-3, and are localized to specific trees along the treeline in site 1 and along the northern side of the creek in site 2, as well as along the powerline in site 3 (where they are frequently observed). American crows (*Corvus brachyrhynchos*) are observed along the treelines in sites 1 and 2, and on the two trees at site 3, but, crucially, not along the powerline, consistent with our field observations that they rarely perch along the powerline. In contrast to these tree-localized species, the swamp sparrow (*Melospiza georgiana*) is predominantly observed on or near the ground away from trees, especially in areas with seasonal ponds. The dickcissel (*Spiza americana*), which is the bird species whose calls we record most frequently, show extreme preference for particular sites and trees (their preference for the more northern of the two trees in front of the house at site 3 matches our subjective observations). The yellow-billed cuckoo (*Coccyzus americanus*) is primarily observed inside the forest, consistent with our experience of it being much more frequently heard than seen.

**Figure 5.**
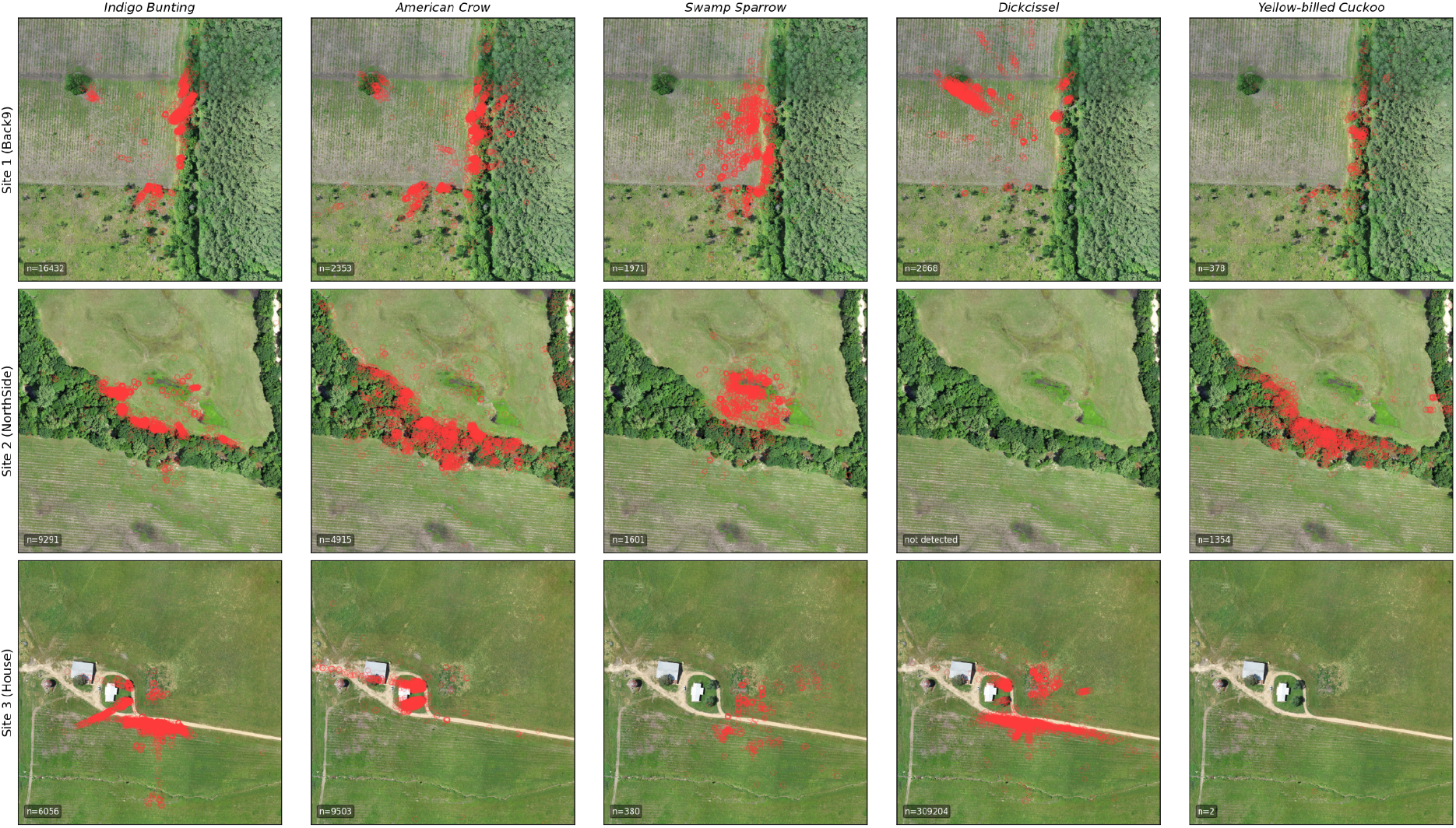
Species-specific spatial distributions of localized calls. Spatial distributions of localized calls for five focal species across the three sites, restricted to events with mean residual TDOA error below 1 ms. Rows are species, columns are sites; each panel spans 300 m centered on the array centroid and is plotted from a random subsample of at most 3,000 points, so density is not comparable across panels. Totals are: indigo bunting (*Passerina cyanea*) 31,852; American crow (*Corvus brachyrhynchos*) 17,099; swamp sparrow (*Melospiza georgiana*) 3,981; dickcissel (*Spiza americana*) 313,820; yellow-billed cuckoo (*Coccyzus americanus*) 1,799. Indigo buntings localize primarily to treelines and powerline perches; American crows localize to trees but rarely to powerlines; swamp sparrows localize near ground-level wetland areas; dickcissels cluster strongly in preferred trees and in open grassland; yellow-billed cuckoos localize predominantly within forest interior. Species are unevenly distributed across sites. For example yellow-billed cuckoo is effectively absent from Site 3 (n = 2) and dickcissel from Site 2 (none detected), so empty panels reflect species occurrence rather than localization failure. These distributions align with known ecological preferences and with field observations at the site.

## Discussion, Limitations and Future Directions

While there are many areas for improvement and further analysis, we conclude that the current system provides robust localization performance under the tested conditions using small arrays of passive recorders in the field. With the introduction of a new generation of inexpensive GPS-synchronized audio recorders (see are currently performing field trials with devices from Frontier Audio, Wildlife Acoustics and AudioMoth), it should be possible to incorporate localization into most soundscape monitoring projects.

As most of our recorders are mounted at ground level, the resulting arrays are largely planar, which is well-known to be non-optimal for localization orthogonal to the plane, which in this case is elevation. The resulting mirror-image ambiguity - a source at height *h* above the array plane produces nearly identical TDOAs to one at *−h* (below the plane) - is trivial to resolve as we do not expect birds to call from inside the Earth. However, the *magnitude* of inferred elevation is poorly constrained in some locations, because the collective intersection geometry of hyperboloids from coplanar recorder pairs provides weak leverage on the vertical coordinate, and thus small TDOA errors translate into large errors in inferred elevation. We therefore treat height estimates as qualitative rather than fully quantitative for now.

There are, however, multiple indications that the elevation estimates are reasonable. Elevations are broadly consistent with the height of landscape: most localizations in unforested areas are near the ground, and in forested areas they are primarily restricted to the height of the trees. There are species-specific differences in height profiles in the same location that are consistent with observations (songbirds in mid-canopy, raptors and crows more elevated, for example). Elevations along the powerline in site 3 are internally consistent and correspond (albeit with significant variance) to the height of the powerlines.

Nonetheless, it is obviously essential to have more precise and accurate elevation measurements, which should be obtainable by adding elevated recorders to the arrays (Dutilleux et al., 2023; Stepanian et al., 2016). Towards this end, we installed elevated (∼7 m) recorders in each array, and results analyzed with these recorders included show reduced vertical uncertainty. However our first generation installations of elevated recorders proved unreliable, and we were thus unable to include these data consistently. Standardizing elevated recorder deployment and integration with ground recorders is a priority for future work.

Overlapping calls and complex acoustic structure can produce ambiguous cross-correlation peaks; while geometric cycle-consistency filtering resolves many such ambiguities, some events remain noisy or poorly constrained, and we have not yet implemented methods to fully identify and flag localization errors introduced by this overlap.

Our current implementation estimates TDOAs using 5-second windows rather than isolating individual call onsets. Manual sub-window annotation improves localization, but we have not yet succeeded in automating this process, so we rely for now on the time-agnostic annotations from BirdNET. This is an area of active investigation elsewhere, and will implement improved call annotation methods as they become available.

Although we are convinced of the high level accuracy of the data by its quantitative characteristics and by the relationship between localizations and the landscape of the sites, an important next step will be the inclusion of ground truth observations localized by observers able to identify and record the precise location and timing of individual calls. This will allow us not only to validate the approach, but to develop improved call-specific error models and quality metrics.

## Code

Code and associated information and outputs are available at https://github.com/cardinal-acoustics/cardinal. The name cardinal is a play on the name of our favorite bird and compass directions. Owing to the historical fandoms of MBE and POB, we do not acknowledge the existence of the baseball team by that name. If you need it to have a backronym, we favor Coordinated Acoustic Recording and Distributed Inference for Naturalistic Animal Localization (CARDINAL).

## Data

The dataset analyzed here is ∼160 Tb of audio files. We are working on a way to make it all available. In the meantime a large sample of regions analyzed here are available at http://carbonranch-cardinal.s3-website.us-east-2.amazonaws.com/.

## AI Use

The work described in the initial version of this preprint was done without the use of any LLM-based AI assistance. For this version Claude (Anthropic) was used to clean up and organize code and figures, to prepare the code for external use, and to efficiently carry out analyses on an expanded dataset. This manuscript was written entirely by MBE.

## Author Contributions

MBE and POB conceived of and planned the project, established the field site, set up the recorder arrays, and travelled to site to collect data cards and perform recorder maintenance. POB designed and built the tone generator used to estimate the speed of sound. MBE and POB collected the drone data used for site imagery. MBE processed the data, wrote all code, carried out all analyses with input from POB. MBE generated all figures and wrote the paper with input from POB. MBE, POB and ASM generated LiDAR data which will be included in a future version of this manuscript to assess localization accuracy, and ASM analyzed these data.

## Acknowledgements

The authors would like to acknowledge the invaluable logistics assistance of Holli Weld in Berkeley, and Nadya Ali and Cody Lockhart in Arkansas.

## Notes

http://carbonranch-cardinal.s3-website.us-east-2.amazonaws.com/

https://github.com/cardinal-acoustics/cardinal

### Competing Interest Statement

The authors have declared no competing interest.

### Summary of Updates

Updated to include a larger input dataset.

### Funding

- Howard Hughes Medical Institute

## References

Aide TM, Corrada-Bravo C, Campos-Cerqueira M, Milan C, Vega G, Alvarez R. 2013. Real-time bioacoustics monitoring and automated species identification. PeerJ 1:e103. doi:10.7717/peerj.103

Cramer O. 1993. The variation of the specific heat ratio and the speed of sound in air with temperature, pressure, humidity, and CO2 concentration. J Acoust Soc Am 93:2510–2516. doi:10.1121/1.405827

Dutilleux G, Sandercock BK, Kålås JA. 2023. Chasing the bird: 3D acoustic tracking of aerial flight displays with a minimal planar microphone array. Bioacoustics 32:622–641. doi:10.1080/09524622.2023.2241420

Gibb R, Browning E, Glover-Kapfer P, Jones KE. 2019. Emerging opportunities and challenges for passive acoustics in ecological assessment and monitoring. Methods in Ecology and Evolution 10:169–185. doi:10.1111/2041-210X.13101

Kahl S, Wood CM, Eibl M, Klinck H. 2021. BirdNET: A deep learning solution for avian diversity monitoring. Ecol Inform 61:101236. doi:10.1016/j.ecoinf.2021.101236

Knapp C, Carter G. 1976. The generalized correlation method for estimation of time delay. IEEE Trans Acoust 24:320–327. doi:10.1109/tassp.1976.1162830

Lapp S, Rhinehart T, Freeland-Haynes L, Khilnani J, Syunkova A, Kitzes J. 2023. OpenSoundscape: An open-source bioacoustics analysis package for Python. Methods Ecol Evol 14:2321–2328. doi:10.1111/2041-210x.14196

Rhinehart TA, Chronister LM, Devlin T, Kitzes J. 2020. Acoustic localization of terrestrial wildlife: Current practices and future opportunities. Ecol Evol 10:6794–6818. doi:10.1002/ece3.6216

Shonfield J, Bayne EM. 2017. Autonomous recording units in avian ecological research: current use and future applications. Avian Conserv Ecol 12. doi:10.5751/ace-00974-120114

Singer A. 2011. Angular synchronization by eigenvectors and semidefinite programming. Appl Comput Harmon Anal 30:20–36. doi:10.1016/j.acha.2010.02.001

Smith J, Abel J. 1987. The spherical interpolation method of source localization. IEEE J Ocean Eng 12:246–252. doi:10.1109/joe.1987.1145217

Stepanian PM, Horton KG, Hille DC, Wainwright CE, Chilson PB, Kelly JF. 2016. Extending bioacoustic monitoring of birds aloft through flight call localization with a three-dimensional microphone array. Ecol Evol 6:7039–7046. doi:10.1002/ece3.2447

Sugai LSM, Silva TSF, Ribeiro JW Jr, Llusia D. 2019. Terrestrial passive acoustic monitoring: Review and perspectives. Bioscience 69:15–25. doi:10.1093/biosci/biy147

Virtanen P, Gommers R, Oliphant TE, Haberland M, Reddy T, Cournapeau D, Burovski E, Peterson P, Weckesser W, Bright J, van der Walt SJ, Brett M, Wilson J, Millman KJ, Mayorov N, Nelson ARJ, Jones E, Kern R, Larson E, Carey CJ, Polat İ, Feng Y, Moore EW, VanderPlas J, Laxalde D, Perktold J, Cimrman R, Henriksen I, Quintero EA, Harris CR, Archibald AM, Ribeiro AH, Pedregosa F, van Mulbregt P, SciPy 1.0 Contributors. 2020. SciPy 1.0: fundamental algorithms for scientific computing in Python. Nat Methods 17:261–272. doi:10.1038/s41592-019-0686-2

Wilson DR, Battiston M, Brzustowski J, Mennill DJ. 2014. Sound Finder: a new software approach for localizing animals recorded with a microphone array. Bioacoustics 23:99–112. doi:10.1080/09524622.2013.827588

